# White matter properties in fronto-parietal tracts predict maladaptive functional activation and deficient response inhibition in ADHD

**DOI:** 10.1101/2023.03.02.530758

**Authors:** Daniel Smullen, Andrew P. Bagshaw, Lilach Shalev, Shlomit Tsafrir, Tamar Kolodny, Carmel Mevorach

## Abstract

Response inhibition, defined as the ability to suppress inappropriate responses, is a key characteristic of adaptive human behaviour. However, in individuals with attention deficit hyperactivity disorder (ADHD) it is often impaired and is linked to broad life outcomes. Previous neuroimaging investigations have indicated a myriad of brain networks in response inhibition, which limit its utility in understanding and overcoming response inhibition difficulties. More recently, it has been suggested that a specific fronto-parietal functional circuitry between the inferior frontal gyrus (IFG) and the intraparietal sulcus (IPS), dictates the recruitment of the IPS in response inhibition in ADHD. To ascertain the critical role of the IFG-IPS functional circuit and its relevance to response inhibition in ADHD, it is crucial to understand the underlying structural architecture of this circuit so that the functional relevance could be interpreted correctly. Here we investigated the white matter pathways connecting the IFG and IPS using seed-based probabilistic tractography on diffusion data in 42 ADHD and 24 neurotypicals and assessed their impact on both the recruitment of IPS in response inhibition scenarios and on response inhibition performance in a Go/No-go task. Our results showed that individual differences in the structural properties of the IPS-IFG circuit, including tract volume and diffusivity, were linked to IPS activation and even predicted response inhibition performance outside the scanner. These findings highlight the structural-functional coupling of the IFG-IPS circuit in response inhibition in ADHD and confirm a structural basis for maladaptive functional top-down control in deficient inhibition in ADHD. Our results also support the notion of ADHD as a continuum and suggest that individual differences in tract-specific functional and structural connectivity could serve as neuromarkers of ADHD.

## 1. Introduction

Inhibiting responses is a survival necessity, from holding our tongue to stay out of trouble to not stepping into the road as a car hurtles round the bend. Nevertheless, impaired response inhibition is one of the most prominent and reproducible behavioural dysfunctions in attention deficit hyperactivity disorder (ADHD, (1–4). Impaired response inhibition may impact quality of life through its contribution to increased risk of school suspension and exclusion (5), reduced job stability (6), increased risk of injury and increased risk of substance abuse (7). Therefore, attenuating the impairment in response inhibition could be pivotal for improving quality of life for individuals with ADHD. A better understanding of the neurobiological mechanisms which underly deficient response inhibition is crucial in order to meet this goal.

A large-scale network of frontal, parietal and striatal brain areas is recruited during response inhibition tasks, including lateral frontal cortex (superior, middle and inferior frontal gyri), the insula, the dorsal medial frontal cortex (supplementary and pre-supplementary motor areas), the anterior cingulate cortex, the inferior parietal cortex, the precuneus, as well as the striatum (8,9). However, the recruitment of many of these areas is not specific to inhibition per-se, but can be attributed to other visual, motor and cognitive demands that these tasks involve. Using a Go/No-go task with a manipulation of target frequency during functional magnetic resonance imaging (fMRI), Kolodny and colleagues (2017) narrowed down these broad frontoparietal activations to specific parietal nodes of the network, which are directly implicated in response inhibition: the bilateral intraparietal sulcus (IPS) and left temporoparietal junction (TPJ) (10). Subsequently, they used the same design to investigate response inhibition in ADHD (11) and found that IPS and TPJ modulation by the inhibitory demand was lacking in ADHD and was mediated by ADHD severity as quantified using the Adult ADHD Self-Report Scale (ASRS; (12–14). In ADHD individuals with mild symptoms, parietal modulation was comparable to controls’, but this modulation dissipated with an increase in ADHD severity. In addition, Kolodny et al. (2020) reported similar effects in task-based functional connectivity between the IPS and the inferior frontal gyrus (IFG). Functional connectivity was modulated by inhibitory load – it was stronger when inhibition was difficult, but this modulation also dissipated with ADHD severity.

Thus, Kolodny and colleagues (10,11) have uncovered a potential mechanism for impaired response inhibition in individuals with ADHD, whereby communication between frontal and parietal nodes is less effective than in neurotypical individuals. This results in reduced top-down modulation of the parietal nodes and consequently reduced sensitivity of these regions to the task context and to inhibitory demand. However, the finding of reduced functional connectivity is not sufficient for us to interpret the mechanism that is underlying it. It could reflect deficiencies in the functional activation of the specific nodes, indicate the use of a different cognitive strategy, or a genuine connectivity deficit that is derived from structural connectivity properties in ADHD.

A structural connectivity deficits as a possible source of this maladaptive functional connectivity would manifest in changes to the underlying structural substrate of the white matter pathways that connect the IFG with the IPS and mediate information transfer between them (15–17). This structural connectivity can be investigated *in-vivo* with diffusion-weighted magnetic resonance imaging (dMRI): the dMRI sequence is sensitive to the random microscopic motion of water molecules, which is constrained by tissue architecture and white matter fibre orientation. Tractography relies on the single-voxel dMRI signal to infer the trajectories of white matter bundles connecting distant regions of the brain, and can be used to assess certain microstructural properties of these bundles (18), including fractional anisotropy (FA), and diffusivity along various axes (axial diffusivity (AD), radial diffusivity (RD) and mean diffusivity (MD)).

Research on white matter in ADHD is somewhat limited and highly inconsistent. In a recent systematic review, Connaughton and colleagues (19) reported that of the twelve studies investigating the corpus callosum in youths with ADHD, eight reported reduced FA and four reported no difference. Similarly, Connaughton and colleagues reported that within the superior longitudinal fasciculus (SLF) there have been findings of both increased and decreased FA, increased and decreased MD and no differences between ADHD and controls. While most research tends to focus on FA and MD, previous findings in ADHD also point to atypicality in global white matter volume (20,21). For this metric too, there is some discrepancy between studies with low global white matter volume in children (Castellanos et al., 2002) and atypically high global white matter volume in adults with ADHD (Seidman et al., 2006), which may point to atypical development of white matter in ADHD. Thus far, research into white matter properties in ADHD has tended to focus primarily on childhood ADHD rather than adults, and on whole-brain white matter or specific cross-hemisphere tracts via the corpus callosum. As such, much remains unknown with respect to white matter atypicality in adults with ADHD. Specifically, how individual differences in white matter properties relate to individual differences in functional brain activation, to behaviour, and indeed to individuals’ clinical phenotypes and symptoms is an underexplored issue that could be crucial to better understand the neural underpinnings of impaired cognitive function in ADHD.

The current work aims to tackle this challenge in the case of response inhibition, utilizing a multi-modal fMRI-dMRI-behavioural dataset. We built upon the previous findings of maladaptive frontoparietal functional connectivity during response inhibition in ADHD to guide a targeted analysis of the dMRI data, using seed-based probabilistic tractography to delineate white matter tracts connecting bilateral IFG and IPS. We first extracted volume and microstructural properties along those specific tracts to test whether any of the structural metrics were atypical in ADHD compared to controls. Then, we assessed if individual differences in the white matter properties of these tracts could predict the individual differences in parietal modulation during inhibitory demand (measured with fMRI). Finally, we also tested if these structural properties could predict performance in a response inhibition task recorded outside the scanner.

## 2. Methods and Materials

### 2.1. Participants

The sample consisted of 42 adults with ADHD and 24 neurotypical controls who also participated in previous fMRI studies by our group (10,11,22). Participants were recruited via advertisements at college and university campuses. All ADHD participants had a previous diagnosis by a qualified clinician and underwent a clinical interview with a psychiatrist who was a member of the research team (author ST) confirming they currently met diagnostic criteria. A history of any non-ADHD neurological or psychiatric condition and non-psychostimulant psychotropic medication use were grounds for exclusion. ADHD current symptom severity was estimated using the total score on the Hebrew version of the Adult ADHD Self-Report Scale (ASRS), a short scale comprised of 18 items corresponding to the DSM diagnostic criteria (13).

Neurotypical participants had no prior history of neurological or psychiatric disorders and no learning disabilities, and were not taking any psychotropic medications. To ensure the absence of attention difficulties, neurotypical participants completed the ASRS, and were included only if they scored within 1 SD of the population’s mean as reported for the Hebrew version (14).

Fourteen participants (ADHD = 9; Control = 5) were excluded following dMRI preprocessing due to poor data quality (see details in *Methods section 2.4.5*). Thus, the final sample consisted of 33 ADHD participants (male = 17) and 19 neurotypical participants (male = 6). A Chi-Square test found that there was no statistically significant association between group and sex (x^2^ (1, N = 52) = 1.942, p = .163). Of the ADHD participants, 1 was medication-naive; 5 had used psychostimulant medication in the past but no longer did; 19 used medication occasionally according to need; and 8 used medication regularly. All participants were required to refrain from using stimulant medication in the 24 hours prior to the experiment. There was no significant difference in white matter metrics between the different medication groups (see *Supplementary Table S1*). All the neurotypical participants were stimulant medication naïve. A breakdown of the demographics for the final sample is shown in *Table 1*, and an equivalent table for the full sample (before exclusions) is provided in the Supplementary Materials (*Table S2*). The study conformed to the Declaration of Helsinki and was approved by the ethics committees of Sheeba Medical Center and of Tel-Aviv University in Israel. All participants provided written informed consent after receiving a complete description of the study.

**Table 1.** Descriptive statistics for demographic and behavioural data

|  | ADHD | Control | t | df | P | Cohen's<br>d |
| --- | --- | --- | --- | --- | --- | --- |
| Sex (M/F) | 17/16 | 6/13 |  |  |  |  |
| Age (years) |  |  | .171 | 50 | .145 | -.049 |
| N | 33 | 19 |  |  |  |  |
| Range (y) | 19-39 | 23-37 |  |  |  |  |
| Mean (y) | 27.5 | 27.7 |  |  |  |  |
| SD (y) | 4.4 | 3.3 |  |  |  |  |
| ASRS Total (Max 90) |  |  | -7.638 | 50 | **<.001\*\*** | 2.200 |
| N | 33 | 19 |  |  |  |  |
| Range | 41-85 | 23-49 |  |  |  |  |
| Mean | 61.5 | 32.2 |  |  |  |  |
| SD | 12.2 | 6.9 |  |  |  |  |
| Response Inhibition Performance<sup>+</sup> |  |  | -1.980 | 43 | **.002\*** | .760 |
| N | 30 | 15 |  |  |  |  |
| Range | 0-.198 | 0-.073 |  |  |  |  |
| Mean | .067 | .033 |  |  |  |  |
| SD | .055 | .016 |  |  |  |  |

<sup>+</sup>Response Inhibition Performance was quantified as the rate of commission errors during the rare-No-go condition in the behavioural version of the Go/No-go task.
Note: \* $p < .05$ , \*\* $p < .001$
Note: SD = standard deviation, df = degrees of freedom
Note: There is variation in the number of participants for each variable due to missing data.

### 2.2. Response inhibition - Go/No-go Task

Response inhibition performance was assessed using a Go/No-go task with a manipulation of target frequency (*Fig. 1*). The task is described in detail by Kolodny and colleagues (10). Briefly, coloured shapes appeared sequentially in the centre of the screen for 100ms each, with varying inter-stimulus intervals (ISI). Participants were instructed to respond quickly via a button click when a red square appeared (Go trial), and withhold response to any other stimuli (No-go trials). The task consisted of two conditions: a rare-No-go condition where 25-30% of the trials were No-go trials) and a prevalent-No-go condition (where 70-75% of trials were No-go trials). The frequent responses in the rare-No-go condition increase the inhibitory demand for No-go trials relative to the prevalent-No-go condition (Kolodny et al., 2017). The Go/No-go task was used both outside the scanner in a separate behavioural session, and during the fMRI session. The task varied slightly between these sessions, in stimulus prevalence (30%/70% in the behavioural session; 25%/75% in the scanner) and in ISI (1-2.5 sec in the behavioural session; 1.8-12 sec in the scanner). Both task versions and the differences between them are described in detail by Kolodny and colleagues (22).

**Figure 1.**
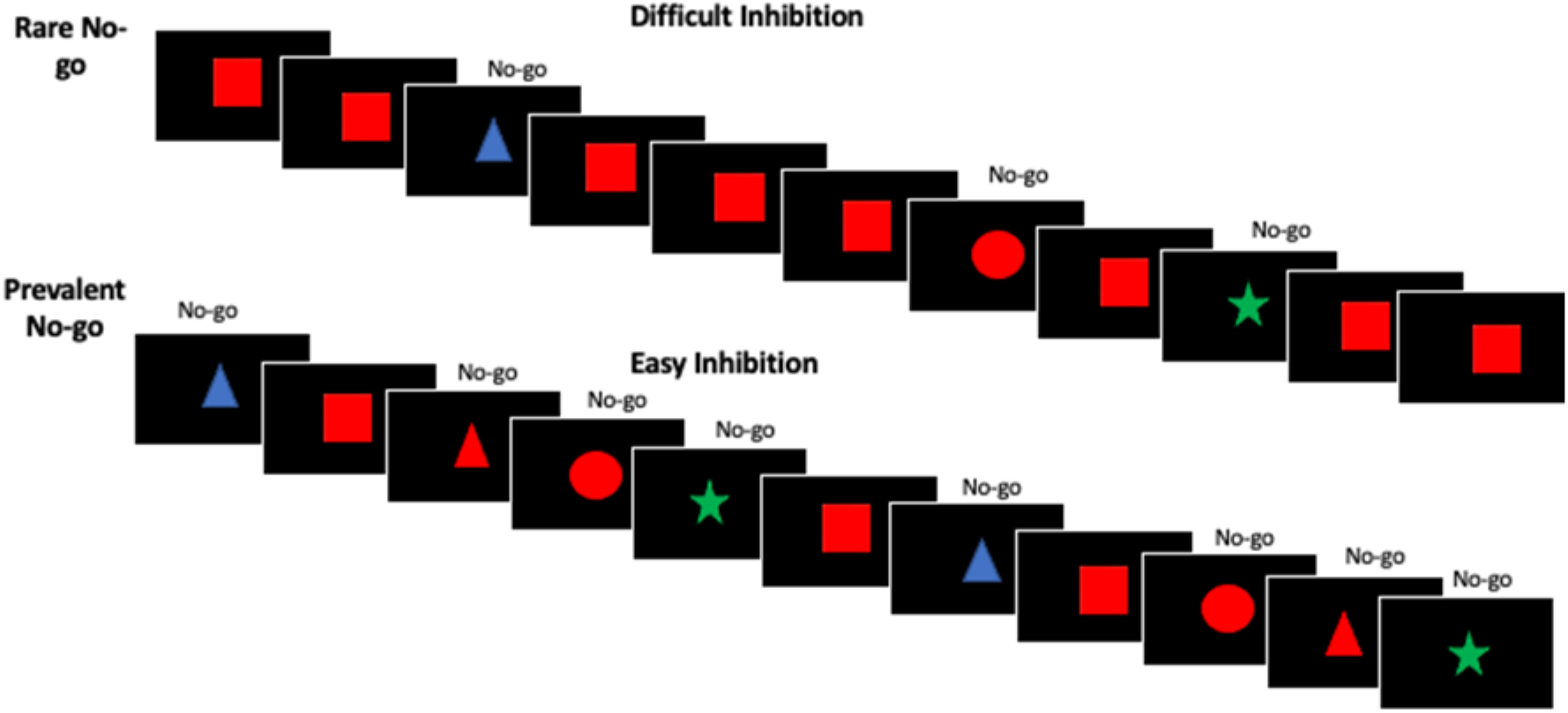
The experimental task, used in the behavioral and fMRI sessions. A series of stimuli were displayed in the centre of the screen. Participants were instructed to respond to a Go stimulus (a red square) using a button click and to ignore all other stimuli (No-go stimuli). There were two task conditions presented in separate runs: rare No-go and prevalent No-go run.

Response inhibition performance was assessed during the behavioural session, that took place prior to the fMRI scanning. Performance was quantified as the percentage of commission errors made in the rare-No-go condition (i.e., the percentage of failed No-go trials, or false alarms. See Table 1 for results), and used as an outcome measure in later regression analyses.

Seven participants (ADHD = 3, Controls = 4) were missing behavioural data, due to technical failure or time constraints during data acquisition.

### 2.3. Neuroimaging

Imaging data were collected in a Siemens 3T Prisma MRI scanner at SCAN@TAU centre in Tel-Aviv University, Israel, using a 64-channel head coil. All data were collected within a single scanning session.

#### 2.3.1 fMRI - parietal modulation during inhibitory demand

fMRI data used in the current research were previously published (Kolodny et al., 2020). Detailed methods including acquisition, task, and processing steps were previously reported, and are therefore only briefly described here. While participants completed the Go/No-go task, 236 functional images were collected using a single-shot 2D gradient-echo echo-planar sequence with the following parameters: slice thickness = 3.6 mm, 33 transverse slices in ascending interleaved order, TR = 2 sec, TE = 35 msec, flip angle = 90, matrix 96 x 96, FOV = 192 mm, for a voxel-wise resolution of 2×2×3.6 mm. An ROI analysis was performed to extract parietal modulation during inhibitory demand: ROIs were defined in right and left IPS and in left TPJ using masks of inhibition-related activation in neurotypicals (masks are available online linked to (10)). In the ADHD group, percent signal change for each condition (rare-No-go and prevalent-No-go) was computed in reference to an implicit baseline and averaged across voxels within each cluster. For each participant and each ROI, the difference between conditions served as the fMRI parietal modulation score, and was used as the outcome measure in the regression models described below.

#### 2.3.2 dMRI Acquisition

dMRI data were acquired using a standard 64-direction protocol using a whole brain echo planar imaging sequence over one phase-encoding direction: TR = 6900ms, TE = 53ms; 76 slices, slice thickness = 1.7mm, no gap; FOV = 197 x 197 mm, matrix size = 216 x 216, providing a cubic resolution of 1.7 x 1.7 x 1.7 mm. 64 diffusion-weighted volumes (b = 1000 sec/mm^2^ and three reference volumes (b = 0 sec/mm^2^) were acquired using a standard diffusion direction matrix.

High-resolution T1-weighted anatomical scans were also collected for each participant to aid co-registration, using a magnetization prepared rapid acquisition gradient echo (MPRAGE) protocol (TR = 1.75 sec, TE = 2.61 msec, T1 = 900 msec, FOV = 220 x 220, matrix = 220 x 220, axial plane, slice thickness = 1 mm, 160 slices, for an isotropic voxel resolution of 1 mm^3^).

#### 2.3.3 dMRI Preprocessing

dMRI preprocessing and processing were performed using the FMRIB Diffusion Toolbox (FDT) in FSL (23–25), following FSL’s standard dMRI processing pipeline. Unweighted (b0) scans were extracted from the dMRI data using the FSL utility tool, fslroi. Brain extraction was performed on one of these images using BET (fractional intensity = 0.5) (26). Eddy-current and motion correction were performed using EDDY and outlier slices were replaced with slices generated by the Gaussian prediction (27,28) as this is suggested to be the most effective method of reducing motion artefacts (29). The number and orientations of crossing fibres within each voxel were modelled using the FDT tool BEDPOSTx. A ball and stick with single diffusion coefficient model was used (30,31).

#### 2.3.4 Quality Assurance

The quality of eddy-current correction was assessed using EDDY QUAD and EDDY SQUAD to flag datasets which potentially contained motion artefacts. Datasets were flagged if their absolute and relative motion metrics were outliers when compared to the control group’s distributions of motion metrics, in order to prevent the potential heightened proneness to in-scanner motion in ADHD masking outliers (32). Flagged datasets were then visually inspected for identifiable motion artefacts. Datasets with a volume containing an identifiable motion artefact were dropped from analysis, as has been previously recommended (33). This resulted in 8 datasets being excluded (ADHD = 6, control = 2). Whole datasets were excluded rather than individually affected volumes as volume removal risks interfering with diffusion metric estimation (34). Furthermore, datasets with more than 20% of volumes containing a replaced slice were considered ‘poor quality’ and also dropped from analysis (ADHD = 3, controls = 3) (35). This resulted in a final dMRI sample of 33 ADHD and 19 controls (*Table 1*).

#### 2.3.5 Probabilistic Tractography

ROIs for seed-based tractography were selected at the right IFG, left IFG, right IPS and left IPS, based on the functional connectivity findings in Kolodny (2017). Seed ROIs were defined using the FIND Atlas (36) and registered to individual subjects’ space using FLIRT (37–39). Probabilistic tractography was performed using Probtrackx (30,31) to produce a 3D structural connectivity distribution for white matter tracts connecting the four ROIs. FSL’s standard settings were used: step length = 0.5mm, samples drawn in each voxel = 5000, subsidiary fibre volume threshold = 0.01, curvature threshold = 0.2. For cross-hemisphere pathways, the corpus callosum was used as a waypoint, defined using the ICBM DTI-81 Atlas (40,41). The end result of probabilistic tractography was connectivity distributions for four tracts (*Fig. 2):* 1) <u>IFG-IFG</u>, a tract connecting the right and left IFG; 2) <u>IPS-IPS</u>, a tract connecting the right and left IPS; 3) <u>Right IFG-IPS</u>, a tract connecting the right IFG and the right IPS; 4) <u>Left IFG-IPS</u>, a tract connecting the left IFG and the left IPS.

**Figure 2.**
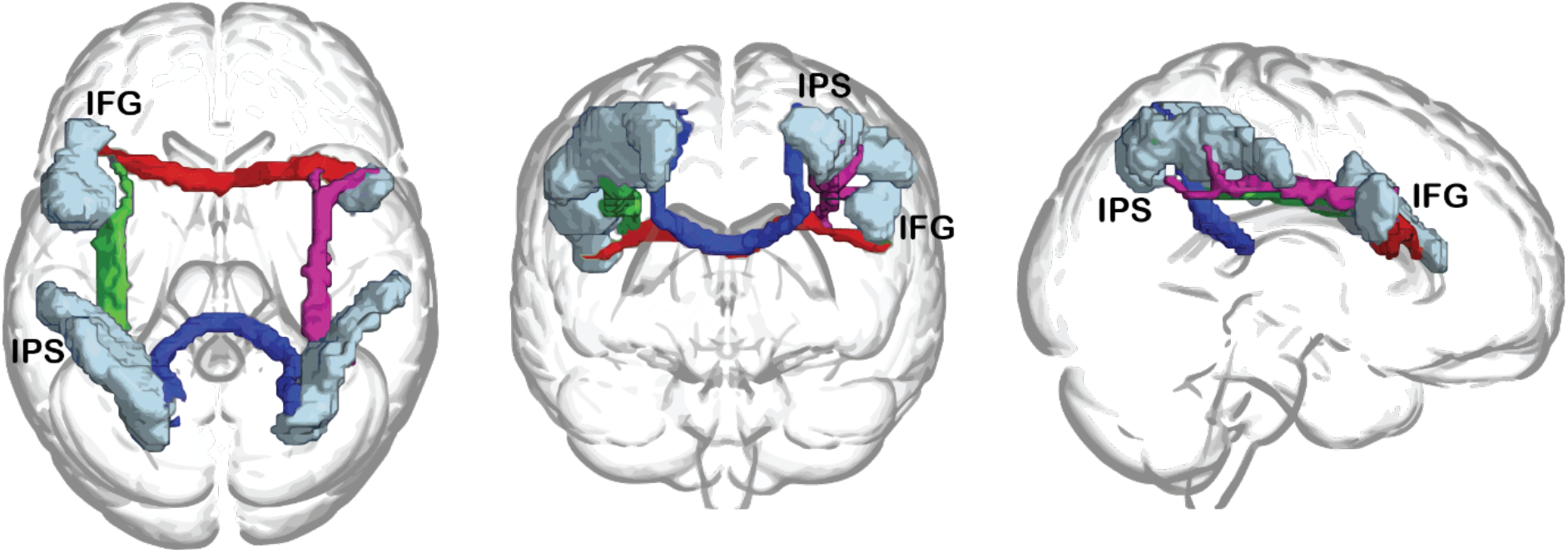
Probabilistic tractography outcomes for an example ADHD participant, warped to standard space and presented in superior, posterior and right views. Seed ROIs in the right and left inferior frontal gyrus (IFG) and intraparietal sulcus (IPS) are shown in light blue. Illustration created by overlaying Probtrackx output on a standard MNI template, using the glass brain setting in MRIcroGL (90).

The IFG-IPS tracts partly overlap with the established Superior Longitudinal Fasciculus (SLF), as determined using the ICBM DTI-81 Atlas. The SLF is a large system of association fibres connecting the frontal and the parietal lobes and prefrontal areas (42–44). The IFG-IFG or IPS-IPS tracts passed through the genu and splenium of the corpus callosum, respectively, but did not clearly overlap with any major white matter pathways from the ICBM DTI-81 Atlas.

#### 2.3.6 Diffusion Tensor Fitting

Diffusion Tensor Fitting was performed at the whole-brain level using DTIFIT to produce FA, AD, RD and MD values in each voxel. Values from voxels contained within the four tracts described above were averaged to produce tract-level metrics.

#### 2.3.7 White matter property extraction

Tract volumes were obtained using fslmaths and fslstats. Spurious tract paths were eliminated prior to volume estimation by thresholding tracts at 15% of maximum path count in each individual and binarizing them to create tract masks (45). To account for the effects of individual brain size differences on tract volume (46), for each participant we determined what percentage of that participant’s whole brain volume was accounted for by a tract’s volume. We then determined what volume this percentage accounted for in MNI standard space. The values produced by DTIFIT from voxels contained within the masks of the four tracts described above were averaged to produce tract-level metrics.

To control for outliers, tracts with within-group volume z > 3 were excluded from subsequent analysis for all metrics, on the basis that this likely represents a poor tract reconstruction. FA, AD, RD and MD outliers were excluded individually using the same criterion (i.e., z > 3). Following outlier detection, all IPS-IPS metrics for one control participant were removed from analysis due to the tract’s volume being an outlier. Individually, one control participant’s IFG-IFG MD and AD were also removed from analysis. For the ADHD group, due to tract volume outliers, all metrics for one participant’s IFG-IFG tract were removed from analysis and all metrics for another participant’s IPS-IPS tract were similarly removed.

#### 2.4 Statistical analysis

Parametric tests were carried out in SPSS 26, whilst non-parametric tests were carried out using custom in-house python-based scripts in Jupyter Notebook (47). To investigate the IFG-IPS structural connectivity, white matter properties for each tract were compared between the ADHD and control groups. Normality of variables was assessed using the Shapiro-Wilk test of normality (48), and group differences were tested using t-tests for normally distributed data, and randomisation tests for data with skewed distributions. Randomisation tests included 100,000 permutations.

To investigate the link between structural connectivity, functional activation and task performance, we used stepwise regression models. The five dMRI metrics (volume, FA, RD, MD & AD) of the four tracts (IPS-IPS, right IPS-IFG, IFG-IFG, left IPS-IFG) were used as potential predictors in these models. Parietal fMRI modulation by inhibitory demand, as well as behavioural response inhibition performance, served as outcome measures. For parietal fMRI modulation, data were available only for the ADHD group, and separate models were used to predict each of the three parietal ROI activations (right IPS, left IPS & left TPJ). To predict behavioural response inhibition task performance, separate models were run for the ADHD and control groups. Thus, five regression models were run in total.

To limit the number of predictors in each regression model and lower the risk of over-fitting, we determined potential predictors to include in the models using Pearson’s correlation coefficients: we first calculated partial correlations between dMRI metrics and the outcome measures, controlling for sex and age. Then, the variables which were correlated with the outcome at a relaxed level of significance (p < 0.10) were included as potential predictors in the corresponding stepwise regression (49). In all stepwise regressions, we controlled for age and sex by including them as potential predictors.

Finally, for all analyses we corrected for multiple comparisons using Benjamini and Hochberg’s Step-Up Procedure (50).

## 3. Results

### 3.1. Group differences in white matter properties

To investigate white matter properties in ADHD, we tested the differences between the ADHD and control groups for the five metrics (volume, FA, MD, AD and RD) in each of the four tracts. Following multiple comparison correction, a randomisation test indicated that AD in the IFG-IFG tract was significantly lower in ADHD (n = 32, mean = 1.25 x 10^-3^ s/mm^2^, sd = 3.80 x 10^-5^) than in controls (n = 18, mean = 1.27 x 10^-3^, sd = 3.22 x 10^-5^; p = .005; *Fig. 3*). It is worth noting that, although they did not reach our pre-defined criterion for exclusion, two particularly low scores from the ADHD group seemed to be driving the effect. Thus, we ran this comparison again with these participants removed, and found that IFG-IFG AD was still significantly lower in the ADHD group (n = 30, mean = 1.25 x 10^-3^ s/mm^2^, sd = 2.82 x 10^-5^) than the control group (n = 18, mean = 1.27 x 10^-3^, sd = 3.22 x 10^-5^; p = .01). No other group differences reached significance (see *Supplementary Table S3*).

**Figure 3.**
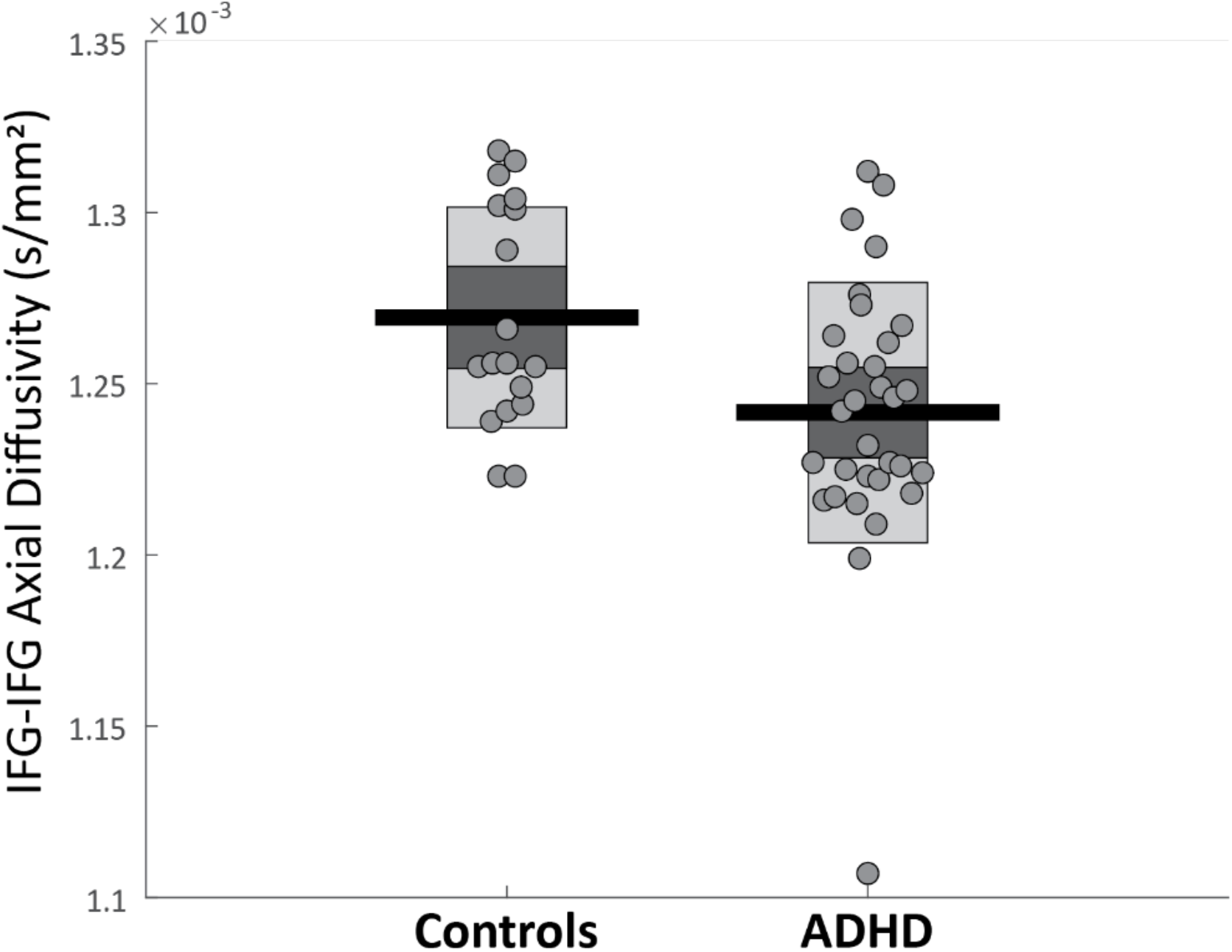
Mean axial diffusivity of the white matter tract connecting the right and left IFG was significantly lower in ADHD than in controls. The group difference remained significant when removing the extreme data point from the ADHD group. Horizontal black lines denote group mean, bars denote 95% confidence intervals, and the central dark area of the bar denotes 1 SD around the mean. Circles represent individual participants.

### 3.2 Predicting parietal functional modulation by inhibitory demand in ADHD from white matter properties

To assess whether structural connectivity in the four tracts could underlie functional activation during inhibitory demand in ADHD, we used stepwise regression models. Potential structural predictors were determined using the procedure outlined in *section 2.4*, resulting in the same four predictors consisting of volume and RD from both the IFG-IFG and left IFG-IPS tracts across the three sites of functional activation. Sex and age were also included in the regressions to control for these variables. Three stepwise regression models were run with activation in the left IPS, left TPJ and right IPS as the outcome measure (separately).

Following multiple comparison correction, all three models were significant, indicating that white matter metrics of the IFG-IFG and left IFG-IPS tracts predicted parietal functional modulation by inhibitory demand. Summaries of these stepwise regression analyses appear below and are visualised in *Fig. 4* (see supplementary *Table S4* for the individual betas for each predictor).

**Figure 4.**
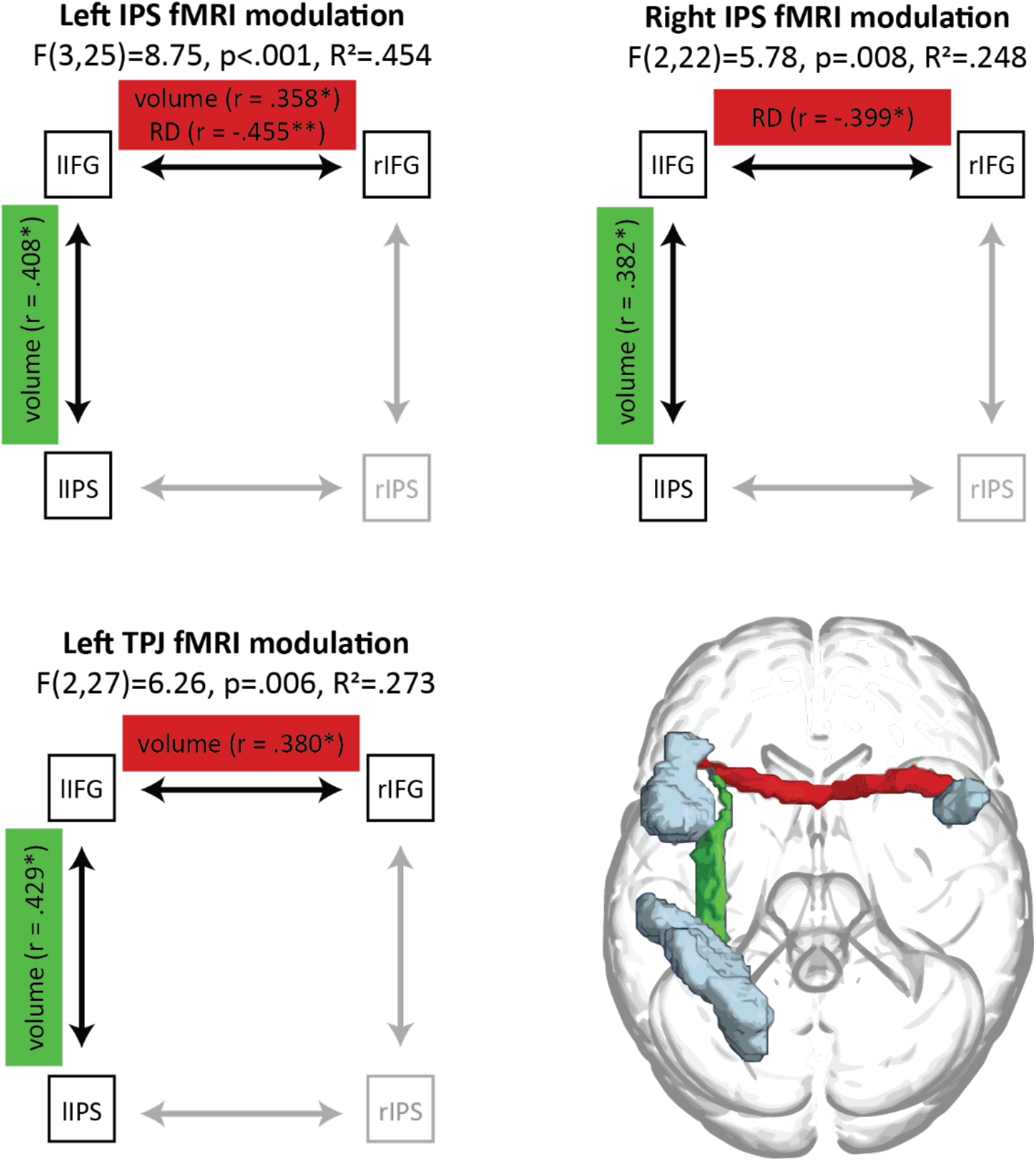
Summary of stepwise regression models predicting fMRI parietal modulations during inhibitory demand in ADHD.

Specifically, the analysis identified a significant model predicting the left IPS functional modulation in the ADHD group (F(3, 25) = 8.750, p < .001, R^2^ = .454). In this model, left IPS modulation was positively predicted by the volumes of the IFG-IFG tract (r = .358, p = .017) and the left IFG-IPS tract (r = .408, p = .007), and negatively predicted by the IFG-IFG tract’s RD (r = −.455, p = .003).

We also found a significant model predicting the right IPS functional modulation in the ADHD group (F(2, 27) = 5.775, p = .008, R^2^ = .248). The left IFG-IPS tract volume again positively predicted right IPS modulation (r = .382, p = .025), and the IFG-IFG tract RD again negatively predicted right IPS modulation (r = −.399, p = .020).

Finally, a significant model was found predicting the left TPJ functional modulation in the ADHD group (F(2,26) = 6.264, p = .006, R^2^ = .273). Here too, volumes of the IFG-IFG tract (r = .380, p = .026) and left IFG-IPS tract (r = .429, p = .013) positively predicted left TPJ modulation.

### 3.3 Predicting response inhibition performance in ADHD and controls from white matter properties

To investigate the ability of white matter properties to predict response inhibition performance in the ADHD and control groups outside the scanner, we again used stepwise regressions. Potential predictors were determined using the procedure outlined in *section 2.4*., and a breakdown of the results can be found in *Supplementary* Table S5. All of the following analyses survived multiple comparison correction and are visualised in *Fig. 5*.

**Figure 5.**
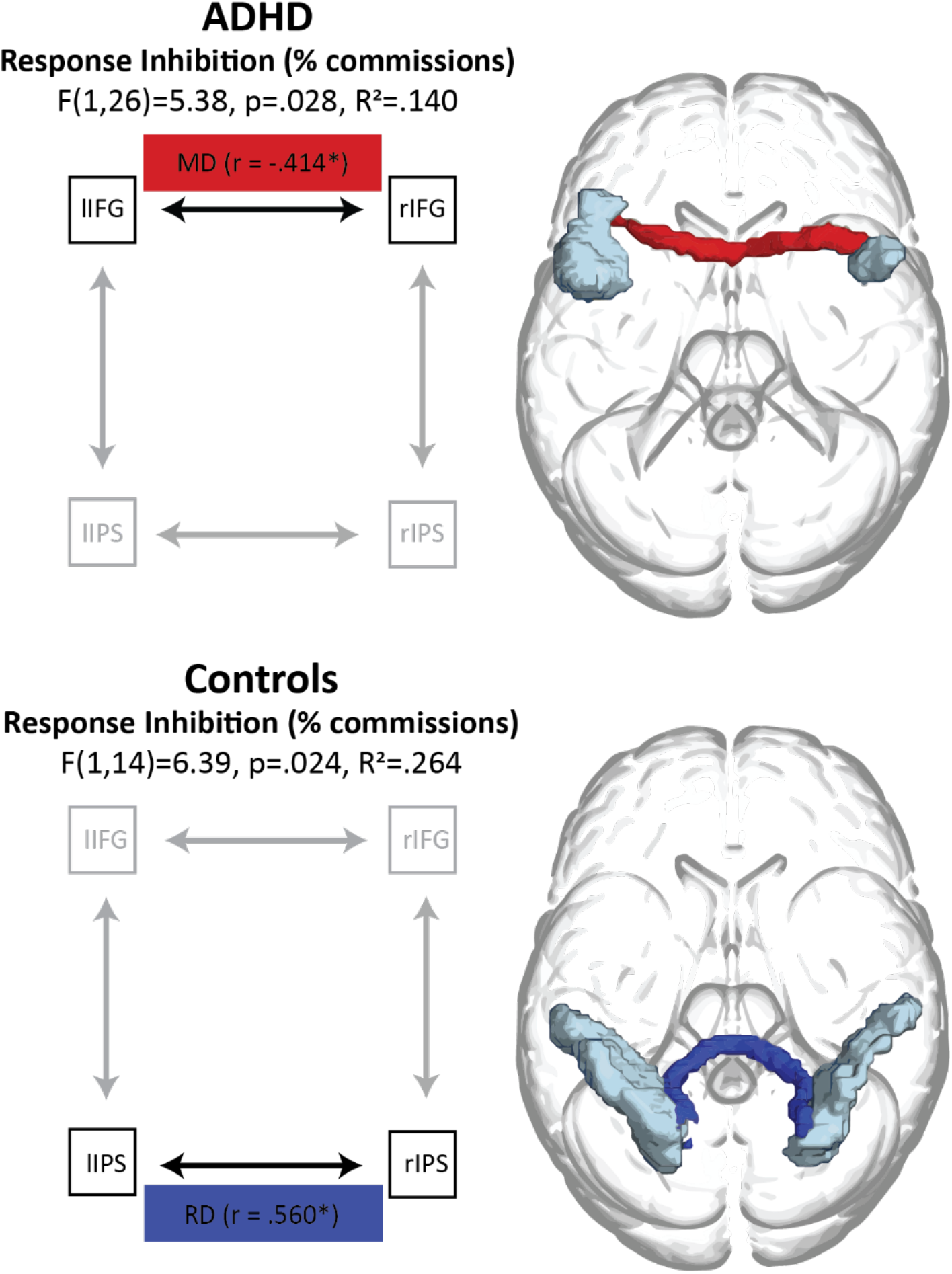
Summary of stepwise regression models predicting response inhibition performance in ADHD and in controls.

Two separate significant models predicted response inhibition performance for the ADHD group and the control group. We found that response inhibition performance was significantly predicted by different structural properties in different tracts for the two groups. In the ADHD group, IFG-IFG tract MD was significantly predictive of response inhibition performance (F(1,26) = 5.380, p = .028, r^2^ = .140), such that higher IFG-IFG tract MD was associated with better response inhibition performance (lower commission errors): r = −.414, p = .028. In controls, response inhibition performance was significantly predicted by IPS-IPS tract RD: F(1, 14) = 6.388, p = .024, R^2^ = .264. The significant contribution made by IPS-IPS tract RD was in a positive direction, with higher IPS-IPS tract RD predicting worse response inhibition performance in controls (higher commission errors): r = .560, p = .024.

### 3.6. Examining potential motion confounds

Although we applied several steps of data cleaning in the dMRI analyses, as detailed in the Methods section, some residual motion artefacts might be present in the data and impact the results (51). As motion is typically higher in ADHD, to ensure structural measures had not been confounded by in-scanner movement we assessed the difference between ADHD and controls’ in-scanner absolute and relative motion metrics. For the final samples, there was no significant difference between the ADHD and control groups’ absolute motion (ADHD: m = .72 mm, sd = .31 mm; controls: m = .62 mm, sd = .16 mm; t(49.76) = 1.523, p = .134). Likewise, there was no significant difference in the relative motion in the two groups (ADHD: m = .35 mm, sd = .03 mm; controls: m = .36 mm, sd = .04 mm; t(50) = .356, p = .724). Thus, it is unlikely that in-scanner motion confounded the current findings. Furthermore, absolute motion and relative motion were not significantly correlated (after multiple comparison correction) with any of the dMRI metrics (see *Supplementary Materials Tables S6 and S7*).

## 4. Discussion

Recent studies from our group (11) pointed to a specific role of a functional circuitry between the IFG and the IPS in deficient response inhibition in ADHD. In the present study we set out to unearth whether this functional activation pattern is coupled with structural connectivity differences in ADHD. This is important because it enables us to interpret the functional differences more meaningfully. If the functional differences previously reported are coupled with structural ones it can point to a systematic organic mechanism that underlies deficient response inhibition in ADHD, and is therefore less likely to results from differences in strategy or. We therefore assessed the relevance of IFG-IPS structural connectivity to the behavioural performance and the functional recruitment of parietal regions for response inhibition in ADHD. Using diffusion MRI and seed-based probabilistic tractography, we assessed the white matter properties of 4 tracts of interest connecting the IFG and the IPS bilaterally. Our findings point to the importance of transcallosal connectivity (volume and diffusivity) between the left and right IFG as well as of a left frontoparietal tract (overlapping with the SLF) in response inhibition in ADHD. On a group level, there were minimal differences between ADHD and controls in the white matter properties (reduced AD) of the IFG-IFG tract. However, on an individual differences’ level within the ADHD group, white matter properties of this tract significantly predicted both the engagement of the parietal regions during task fMRI (volume and RD) and behavioural performance in a response inhibition task (MD). White matter properties of the left fronto-parietal tract (volume) also significantly predicted the engagement of the parietal cortex during the response inhibition task in this group. In contrast to individuals with ADHD, in control participants white matter properties of the posterior IPS-IPS tract (RD) significantly predicted response inhibition performance.

Some evidence of a role for transcallosal IFG to IFG connectivity in ADHD was provided by the reduced AD in ADHD compared to controls. Reduced AD has been proposed to reflect reduced calibre of, or injury to, axons, or reduced coherence in axon orientation (52). However, the anatomy which underpins AD and thus the implication of reduced AD is debated (53). Our findings of atypical IFG-IFG AD in ADHD suggests that this tract is compromised in the disorder, which is in line with reports that the IFG is a key node for cognitive and attentional control, whose activity is impaired in ADHD (54–56). Our findings support this notion in highlighting the importance of this tract in ADHD generally, although with the caveat that the group differences in AD were not large and possibly driven by a small number of participants.

Perhaps more informatively, our results also implicate the IFG-IFG tract’s structure specifically in the context of response inhibition. We found that increased IFG-IFG RD, which reflects water molecule diffusivity magnitude perpendicular to fibre tracts, partly predicted reduced left and right IPS modulation during inhibitory demand as measured with fMRI. As with AD, the anatomical underpinnings of RD are debated, but increased RD has been proposed to reflect a loss of myelin, loss of axons, or reduced axonal packing density (52). In addition to the tract’s RD, our results also implicated tract volume, with increased IFG-IFG tract volume predicting greater left IPS and left TPJ fMRI modulation during inhibitory demand (i.e., modulation that is more similar to what is observed in control participants). Factors proposed to drive an alteration in white matter volume include number of axons, thickness of myelin around axons, number of axons with a myelin sheath, axonal branching and axonal crossing (57).

Importantly, IFG-IFG white matter properties not only predicted functional engagement of parietal nodes for increased inhibitory demand, but also behavioural response inhibition performance in a Go/No-go task among participants with ADHD. Increased IFG-IFG tract MD (the average of AD and RD) predicted better response inhibition performance. Notably, this was not the case for the control group. Instead, their response inhibition performance was predicted by the RD of the parietal tract connecting the left and right IPS. These differential results suggest that response inhibition may be executed differently in the brain of individuals with and without ADHD, and rely more heavily on certain circuitry in each group. Specifically, the IFG is associated with cognitive and attentional control (58) but also involved in a broad range of other cognitive functions, and has been suggested as one of the key domain-general areas in the brain (59). The fact that individual differences in the IFG-IFG tract in ADHD predicts behavioural performance may suggest that their difficulty in response inhibition stems from an alteration in a more general cognitive function (e.g. context monitoring), or that during response inhibition they recruit a broader neural network instead of specialized circuits, which may be less efficient.

The IFG-IFG tract in the current study crosses between the hemispheres through the anterior part of the corpus callosum. The findings we report here are consistent with previous reports highlighting reduced corpus callosum volume in children with ADHD, and its association with reduced sustained attention and attentional control (60,61). AD in the corpus callosum has also been implicated in ADHD, with increased AD linked with greater symptom severity, and poorer attention and working memory (62). Thus, the current findings of atypical IFG-IFG connectivity in the more severe ADHD participants may be related to general corpus callosum atypicalities in ADHD and their effect on interhemispheric communication. However, as the interhemispheric parietal tract was not implicated in ADHD by the current research, it may point to a more specific atypicality in the frontal tract or in the genu of the corpus callosum through which this tract passes. Future research using spatial tract profiling and corpus callosum segmentation is required to further clarify this issue.

Apart from the IFG-IFG tract, the left hemisphere fronto-parietal tract connecting the IFG and the IPS was also related to response inhibition functional modulation in ADHD. We found that increased left IFG-IPS tract volume predicted increased activity in the left IPS, right IPS, and left TPJ during inhibitory demand. This further supports the notion that IFG-IPS connectivity is important in ADHD for engaging parietal cortices when the response inhibition demand increases (11). Using intracranial EEG and direct electrical stimulation (DES), it has been found that critical time windows for the IPS and the IFG during response inhibition are highly similar, and thus it has been hypothesized that IFG and IPS processes run separately but in parallel during inhibitory demand (63–65). An alternative hypothesis proposes that the two regions do not behave separately, but rather that the inferior frontal cortex inhibits down-stream parietal activity during inhibitory demand (66). Our finding implicating IFG-IPS communication in ADHD-impaired response inhibition suggests that this communicative pathway is important for successful response inhibition, lending support to the latter theory that the inferior frontal cortex modulates down-stream parietal activity during inhibitory demand.

The IFG-IPS tract makes up, in part, the ventrolateral subdivision of the superior longitudinal fasciculus (SLF III). Structure-function relationships between the SLF and specific cognitive functions are yet to be fully elucidated, but associations have been suggested with perceptual organization and working memory (67), sustained attention (68), and importantly for the interpretation of current results, response inhibition (69). Indeed, previous research has also implicated white matter properties of the SLF in children and in adults with ADHD (70–72). It is worth noting that our findings suggest the left tract, but not the right tract, to be important in response inhibition. This may be surprising given the central role of the right IFG, specifically, in previous studies of response inhibition (73,74). However, since our findings also demonstrate a role for the IFG-IFG tract, they do not detract from the involvement of the right IFG in the inhibition-related circuitry. Furthermore, our findings are consistent with a handful of studies highlighting the role of the left IFG in inhibition (75,76)and studies indicating that its integrity is critical for successful implementation of response inhibition (77).

In the current discussion we have implicitly considered the link between white matter structure and response inhibition to be of a nature whereby structural metrics underlie functional activation and subsequently cause response inhibition performance variation. However, the structure-function relationship is known to be bidirectional, and training has been shown to induce changes in white matter properties (57). For example, increased FA in somatosensory and visual cortices followed tactile braille reading training (78), increased FA in the right IPS followed juggling training (79), and reduced MD in the left arcuate fasciculus followed intensive reading intervention (80). Thus, the findings of the current study, highlighting individual differences in structural and functional connectivity in the IPS-IFG circuit as potential neuromarkers of response inhibition, suggest that the IFG-IFG and IPS-IFG white matter tracts could be used as potential targets to study intervention outcomes. Future research may investigate the plasticity of this neural circuit, perhaps via training.

Several training regimes have been demonstrated to improve executive functioning and inhibitory control, including computerized cognitive training (81), mindfulness practice (82) and neurofeedback (83), but none of the existing studies tracked induced changes in structural brain properties.

Apart from training, changes to white matter are also triggered by medications (84). Specifically, modest increases in FA have been shown to follow methylphenidate treatment in boys with ADHD (85). It is possible that other factors, including nutrition (86) and sleep patterns (86) could also invoke white matter plasticity. These provide further potential avenues to explore in order to test whether the IPS-IFG structural properties are plastic, and whether changes to this underlying white matter circuit could lead to changes in function and behaviour. Importantly, deficient response inhibition is impactful throughout the entire life span. Thus, it is important for future studies to investigate the emergence of white matter differences in ADHD throughout development, and the potential of early intervention to shape white matter connectivity development from a young age.

There are a number of limitations to the current study. First, dMRI data was collected using relatively basic acquisition parameters: a single b-value and a single phase-encoding direction. A single b-value limits the robustness of cross-fibre modelling, relative to current state-of-the-art multi-shell acquisitions. As no opposite phase encoding direction scan was acquired, motion correction could not include an estimation of motion’s impact on susceptibility-induced distortions. Whilst this step is not mandatory, it is recommended and could be specifically important when one group of participants is more prone for motion, as is the case for ADHD participants (87–89). We addressed possible motion artefacts by strict preprocessing quality control and excluded data flagged for motion. We further showed that in the retained data, there was no significant group difference in in-scanner motion, and motion parameters did not correlate with any white matter metrics. Thus, although distortion correction could have been beneficial, we believe the current findings are not driven by motion artefacts.

The current study was also limited in regard to data availability. As the parietal ROIs used to extract fMRI activation during response inhibition were defined by Kolodny and colleagues (10) using the same control participants included in the current study, we opted not to use their fMRI data as dependent variables in regression models, to avoid circularity. Thus, we could only investigate the role of IFG-IPS structural connectivity in parietal modulation by inhibitory demand in ADHD, but not in controls. Collecting data in a novel group of control participants will enable to test whether the same white matter tracts play a similar role in the two populations.

To conclude, the current study demonstrated a role for white matter structure in the atypical recruitment of fronto-parietal networks in ADHD during response inhibition and as such provided evidence for the coupling of structural and functional connectivity between the IFG and IPS in response inhibition in ADHD. Indeed, microstructural properties of the cross-hemispheric IFG-IFG tract and the left fronto-parietal IFG-IPS tract predict not only fMRI activation but also response inhibition performance in ADHD, as measured behaviourally in the lab (outside the scanner). Together, these findings highlight deficient top-down control brought about by maladaptive connectivity as a mechanism of inhibition difficulty in ADHD. Our results point to the potential of individual differences in tract-specific functional and structural connectivity properties as neuromarkers of ADHD and as loci for future intervention programs.

## Acknowledgments

This study was funded by grant no. 3-7331 from the Chief Scientist of the Israeli Ministry of Health to author LS. We thank Pnina Stern for assistance with participant recruitment and behavioural data collection, and Maya Ankaoua and Natalie Biderman for assistance with imaging data collection.

