## Supplementary Materials for "White matter properties in fronto-parietal tracts predict maladaptive functional activation and deficient response inhibition in ADHD"

| **Table S1.** ANOVA results investigating the difference in white matter metrics between  medication usage groups due to medication’s known effect on white matter. Whilst tests  were limited by uneven group sizes and no medication-naïve group, we nevertheless found  that medication had no significant effect on white matter properties. | | | | | |
| --- | --- | --- | --- | --- | --- |
|  | n | df | Mean | F | p |
| lIFG-lIPS Volume |  | 3, 29 |  | .076 | .972 |
| Past Medication Use | 5 |  | 2865.73 |  |  |
| Occasional Medication Use | 19 |  | 3145.15 |  |  |
| Regular Medication Use | 8 |  | 3093.07 |  |  |
| rIFG-rIPS Volume |  | 3, 29 |  | .5 | .685 |
| Past Medication Use | 5 |  | 2463.11 |  |  |
| Occasional Medication Use | 19 |  | 2697.35 |  |  |
| Regular Medication Use | 8 |  | 2784.12 |  |  |
| IFG-IFG Volume |  | 3, 28 |  | .871 | .468 |
| Past Medication Use | 5 |  | 3600.63 |  |  |
| Occasional Medication Use | 18 |  | 3666.50 |  |  |
| Regular Medication Use | 8 |  | 3495.14 |  |  |
| IPS-IPS Volume |  | 3, 28 |  | .966 | .422 |
| Past Medication Use | 5 |  | 3443.78 |  |  |
| Occasional Medication Use | 19 |  | 4086.29 |  |  |
| Regular Medication Use | 7 |  | 3747.73 |  |  |
| lIFG-lIPS FA |  | 3, 29 |  | .553 | .650 |
| Past Medication Use | 5 |  | .484 |  |  |
| Occasional Medication Use | 19 |  | .492 |  |  |
| Regular Medication Use | 8 |  | .501 |  |  |
| rIFG-rIPS FA |  | 3, 29 |  | .174 | .913 |
| Past Medication Use | 5 |  | .487 |  |  |
| Occasional Medication Use | 19 |  | .488 |  |  |
| Regular Medication Use | 8 |  | .489 |  |  |
| IFG-IFG FA |  | 3, 28 |  | .737 | .593 |
| Past Medication Use | 5 |  | .476 |  |  |
| Occasional Medication Use | 18 |  | .494 |  |  |
| Regular Medication Use | 8 |  | .481 |  |  |
| IPS-IPS FA |  | 3, 28 |  | .729 | .543 |
| Past Medication Use | 5 |  | .576 |  |  |
| Occasional Medication Use | 19 |  | .568 |  |  |
| Regular Medication Use | 7 |  | .582 |  |  |
| lIFG-lIPS MD |  | 3, 29 |  | .364 | .779 |
| Past Medication Use | 5 |  | .0703 x 10^-3^ |  |  |
| Occasional Medication Use | 19 |  | .0708 x 10^-3^ |  |  |
| Regular Medication Use | 8 |  | .0708 x 10^-3^ |  |  |
| rIFG-rIPS MD |  | 3, 29 |  | .849 | .479 |
| Past Medication Use | 5 |  | .0717 x 10^-3^ |  |  |
| Occasional Medication Use | 19 |  | .0724 x 10^-3^ |  |  |
| Regular Medication Use | 8 |  | .0718 x 10^-3^ |  |  |
| IFG-IFG MD |  | 3, 28 |  | .746 | .534 |
| Past Medication Use | 5 |  | .0765 x 10^-3^ |  |  |
| Occasional Medication Use | 18 |  | .0771 x 10^-3^ |  |  |
| Regular Medication Use | 8 |  | .0773 x 10^-3^ |  |  |
| IPS-IPS MD |  | 3, 28 |  | 1.342 | .281 |
| Past Medication Use | 5 |  | .0765 x 10^-3^ |  |  |
| Occasional Medication Use | 19 |  | .0774 x 10^-3^ |  |  |
| Regular Medication Use | 7 |  | .0765 x 10^-3^ |  |  |
| lIFG-lIPS AD |  | 3, 29 |  | 1.871 | .157 |
| Past Medication Use | 5 |  | 1.10 x 10^-3^ |  |  |
| Occasional Medication Use | 19 |  | 1.14 x 10^-3^ |  |  |
| Regular Medication Use | 8 |  | 1.12 x 10^-3^ |  |  |
| rIFG-rIPS AD |  | 3, 29 |  | .306 | .821 |
| Past Medication Use | 15 |  | 1.11 x 10^-3^ |  |  |
| Occasional Medication Use | 19 |  | 1.11x 10^-3^ |  |  |
| Regular Medication Use | 8 |  | 1.11 x 10^-3^ |  |  |
| IFG-IFG AD |  | 3, 28 |  | .280 | .840 |
| Past Medication Use | 4 |  | 1.12 x 10^-3^ |  |  |
| Occasional Medication Use | 19 |  | 1.12 x 10^-3^ |  |  |
| Regular Medication Use | 8 |  | 1.12 x 10^-3^ |  |  |
| IPS-IPS AD |  | 3, 29 |  | .051 | .985 |
| Past Medication Use | 5 |  | 1.13 x 10^-3^ |  |  |
| Occasional Medication Use | 19 |  | 1.14 x 10^-3^ |  |  |
| Regular Medication Use | 8 |  | 1.13 x 10^-3^ |  |  |
| lIFG-lIPS RD |  | 3, 29 |  | .093 | .963 |
| Past Medication Use | 5 |  | 0.504 x 10^-3^ |  |  |
| Occasional Medication Use | 19 |  | 0.501 x 10^-3^ |  |  |
| Regular Medication Use | 8 |  | 0.498 x 10^-3^ |  |  |
| rIFG-rIPS RD |  | 3, 29 |  | .838 | .484 |
| Past Medication Use | 5 |  | 0.510 x 10^-3^ |  |  |
| Occasional Medication Use | 19 |  | 0.516 x 10^-3^ |  |  |
| Regular Medication Use | 8 |  | 0.515 x 10^-3^ |  |  |
| IFG-IFG RD |  | 3, 29 |  | .448 | .720 |
| Past Medication Use | 5 |  | 0.540 x 10^-3^ |  |  |
| Occasional Medication Use | 19 |  | 0.539 x 10^-3^ |  |  |
| Regular Medication Use | 8 |  | 0.545 x 10^-3^ |  |  |
| IPS-IPS RD |  | 3, 29 |  | .2.293 | .099 |
| Past Medication Use | 5 |  | 0.473 x 10^-3^ |  |  |
| Occasional Medication Use | 19 |  | 0.485 x 10^-3^ |  |  |
| Regular Medication Use | 8 |  | 0.467 x 10^-3^ |  |  |

**Table S2.** Descriptive statistics for demographic and behavioural data prior to outlier removal

|  | ADHD | Control | t | df | P | Cohen’s d |
| --- | --- | --- | --- | --- | --- | --- |
| Sex (M/F) | 20/23 | 8/16 |  |  |  |  |
| Age (years) |  |  | -.537 | 65 | .595 | -.137 |
| N | 43 | 24 |  |  |  |  |
| Range (y) | 19-39 | 19-38 |  |  |  |  |
| Mean (y) | 26.7 | 27.3 |  |  |  |  |
| SD (y) | 4.4 | 4.2 |  |  |  |  |
| ASRS Total (Max 90) |  |  | 9.813 | 65 | **.005*** | 2.500 |
| N | 43 | 24 |  |  |  |  |
| Range | 41-85 | 23-49 |  |  |  |  |
| Mean | 63.0 | 37.7 |  |  |  |  |
| SD | 11.6 | 6.7 |  |  |  |  |
| Response Inhibition Performance^+^ |  |  | 2.744 | 51 | **<.001**** | .786 |
| N | 34 | 19 |  |  |  |  |
| Range | 0-.198 | 0-.073 |  |  |  |  |
| Mean | .064 | .028 |  |  |  |  |
| SD | .053 | .016 |  |  |  |  |
| ^+^Response Inhibition Performance was quantified as the rate of commission errors during the rare-No-go condition in the behavioural version of the Go/No-go task.  Note: * p < .05, ** p < .001  Note: SD = standard deviation, df = degrees of freedom  Note: There is variation in the number of participants for each variable due to missing data. | | | | | | |

**Table S3.** A summary of t-tests and randomisation tests assessing the difference in white matter properties between the ADHD and control groups which were either non-significant or did not survive multiple comparison correction

|  | ADHD | Control | t | df | P | |
| --- | --- | --- | --- | --- | --- | --- |
| lIFG-lIPS volume |  |  | 1.243 | 50 | | .219 |
| N | 33 | 19 |  |  |  | |
| Mean (mm^3^) | 3082.34 | 2679.89 |  |  |  | |
| SD (mm^3^) | 1180.29 | 1015.77 |  |  |  | |
| lIFG-lIPS FA |  |  | -.361 | 50 | .167 | |
| N | 33 | 19 |  |  |  | |
| Mean | 4.93 x 10^-1^ | 4.95 x 10^-1^ |  |  |  | |
| SD | 2.40 x 10^-2^ | 2.30 x 10^-2^ |  |  |  | |
| lIFG-lIPS MD |  |  | -1.465 | 50 | .237 | |
| N | 33 | 19 |  |  |  | |
| Mean | 7.08 x 10^-4^ | 7.18 x 10^-4^ |  |  |  | |
| SD | 1.47 x 10^-5^ | 3.31 x 10^-5^ |  |  |  |  |
| lIFG-lIPS AD |  |  | -1.608 | 50 | .114 | |
| N | 33 | 19 |  |  |  | |
| Mean | 1.12 x 10^-3^ | 1.14 x 10^-3^ |  |  |  | |
| SD | 2.90 x 10^-5^ | 4.35 x 10^-5^ |  |  |  | |
| lIFG-lIPS RD |  |  | -.890 | 50 | .445 | |
| N | 33 | 19 |  |  |  | |
| Mean | 5.00 x 10^-4^ | 5.07 x 10^-4^ |  |  |  | |
| SD | 2.00 x 10^-5^ | 3.43 x 10^-5^ |  |  |  | |
| rIFG-rIPS volume |  |  |  |  | .077 | |
| N | 33 | 19 |  |  |  | |
| Mean (mm^3^) | 2748.12 | 1008.81 |  |  |  | |
| SD (mm^3^) | 2387.73 | 531.97 |  |  |  | |
| rIFG-rIPS FA |  |  | -1.166 | 50 | .249 | |
| N | 33 | 19 |  |  |  | |
| Mean | 4.87 x 10^-1^ | 4.95 x 10^-1^ |  |  |  | |
| SD | 2.23 x 10^-2^ | 2.04 x 10^-2^ |  |  |  | |
| rIFG-rIPS MD |  |  | -.043 | 50 | .966 | |
| N | 33 | 19 |  |  |  | |
| Mean | 7.22 x 10^-4^ | 7.22 x 10^-4^ |  |  |  | |
| SD | 2.14 x 10^-5^ | 2.58 x 10^-5^ |  |  |  | |
| rIFG-rIPS AD |  |  | -.699 | 50 | .487 | |
| N | 33 | 19 |  |  |  | |
| Mean | 1.14 x 10^-3^ | 1.14 x 10^-3^ |  |  |  | |
| SD | 3.28 x 10^-5^ | 4.24 x 10^-5^ |  |  |  | |
| rIFG-rIPS RD |  |  | .477 | 50 | .635 | |
| N | 33 | 19 |  |  |  | |
| Mean | 5.15 x 10^-4^ | 5.11 x 10^-4^ |  |  |  | |
| SD | 2.47 x 10^-5^ | 2.45 x 10^-5^ |  |  |  | |
| lIFG-rIFG volume |  |  |  |  | .086 | |
| N | 32 | 19 |  |  |  | |
| Mean (mm^3^) | 3695.83 | 3067.17 |  |  |  | |
| SD (mm^3^) | 1649.05 | 1480.51 |  |  |  | |
| lIFG-rIFG FA |  |  |  |  | .083 | |
| N | 32 | 19 |  |  |  | |
| Mean | 4.87 x 10^-1^ | 5.00 x 10^-1^ |  |  |  | |
| SD | 3.03 x 10^-2^ | 3.27 x 10^-2^ |  |  |  | |
| lIFG-rIFG MD |  |  | -.723 | 48 | .473 | |
| N | 32 | 18 |  |  |  | |
| Mean | 7.72 x 10^-4^ | 7.77 x 10^-4^ |  |  |  | |
| SD | 2.37 x 10^-5^ | 1.74 x 10^-5^ |  |  |  | |
| lIFG-rIFG RD |  |  | 1.222 | 49 | .228 | |
| N | 32 | 19 |  |  |  | |
| Mean | 5.42 x 10^-4^ | 5.30 x 10^-4^ |  |  |  | |
| SD | 3.57 x 10^-5^ | 3.18 x 10^-5^ |  |  |  | |
| lIPS-rIPS volume |  |  |  |  | .021 | |
| N | 32 | 18 |  |  |  | |
| Mean (mm^3^) | 3863.55 | 1122.91 |  |  |  | |
| SD (mm^3^) | 4842.95 | 2109.02 |  |  |  | |
| lIPS-rIPS FA |  |  | .260 | 48 | .796 | |
| N | 32 | 18 |  |  |  | |
| Mean | 5.71 x 10^-1^ | 5.70 x 10^-1^ |  |  |  | |
| SD | 2.83 x 10^-2^ | 2.75 x 10^-2^ |  |  |  | |
| lIPS-rIPS MD |  |  | -1.571 | 47 | .123 | |
| N | 32 | 17 |  |  |  | |
| Mean | 7.72 x 10^-4^ | 7.85 x 10^-4^ |  |  |  | |
| SD | 2.60 x 10^-5^ | 2.90 x 10^-5^ |  |  |  | |
| lIPS-rIPS AD |  |  | -.962 | 48 | .341 | |
| N | 32 | 18 |  |  |  | |
| Mean | 1.35 x 10^-3^ | 1.37 x 10^-3^ |  |  |  | |
| SD | 6.32 x 10^-5^ | 5.306 x 10^-5^ |  |  |  | |
| lIPS-rIPS RD |  |  | -1.055 | 48 | .297 | |
| N | 32 | 18 |  |  |  | |
| Mean | 4.81 x 10^-4^ | 4.90 x 10^-4^ |  |  |  | |
| SD | 2.96 x 10^-5^ | 3.61 x 10^-5^ |  |  |  | |
| **Note:** All P values are raw  **Note:** Tests without t values were randomisation tests | | | | | | |

| **Table S4.** A summary of the stepwise regression model predicting left IPS, right IPS and left TPJ modulations during inhibitory demand by white matter potential predictors within the ADHD group. | | | |
| --- | --- | --- | --- |
|  | r | β | t |
| lIPS modulation (R^2^ = .454) |  |  |  |
| Age | .288 | .147 | .956 |
| Sex | .120 | -.544 | -.954 |
| rIFG-lIFG volume | .382** | .358** | 2.555** |
| rIFG-lIFG RD | -.471*** | -.455*** | -3.252*** |
| lIFG-lIPS volume | .395*** | .408** | 2.920** |
| lIFG-lIPS RD | -.043 | -- | -- |
| rIPS modulation (R^2^ = .248) |  |  |  |
| Age | .077 | -.021 | -.122 |
| Sex | .117 | -.098 | -.543 |
| rIFG-lIFG volume | .324 | -- | -- |
| rIFG-lIFG RD | -.392** | -.399** | -2.476** |
| lIFG-lIPS volume | .375** | .382** | 2.374** |
| lIFG-lIPS RD | -.138 | -- | -- |
| lTPJ modulation (R^2^ = .273) |  |  |  |
| Age | .288 | .136 | .787 |
| Sex | .257 | .091 | .497 |
| rIFG-lIFG volume | .375** | .380** | 2.360** |
| rIFG-lIFG RD | -.268 | -- | -- |
| lIFG-lIPS volume | .425** | .429** | 2.666** |
| lIFG-lIPS RD | -.065 | -- | -- |
| Note: * p < .01, ** p < .05, *** p < .005 | | | |

| **Table S5.** A summary of the stepwise regression model predicting response inhibition performance by white matter potential predictors within the ADHD and control groups. | | | |
| --- | --- | --- | --- |
|  | r | β | T |
| ADHD Response Inhibition (R^2^ = .140) |  |  |  |
| Age | -.044 | -.136 | -.742 |
| Sex | -.271 | .165 | .916 |
| rIFG-lIFG MD | -.406** | -.414** | -2.319** |
| rIFG-lIFG RD | -.331** | .190 | -.648 |
| lIFG-lIPS FA | .337 | .259 | 1.484 |
| rIFG-rIPS MD | -.383** | -.281 | -1.591 |
| rIFG-rIPS RD | -.414** | -.298 | -1.692 |
| lIPS-rIPS RD | .125 | -- | -- |
| Controls Response Inhibition (R^2^ = .264) |  |  |  |
| Age | -.367 | -.265 | 1.189 |
| Sex | .439* | .362 | 1.726 |
| rIFG-lIFG MD | -.089 | -- | -- |
| rIFG-lIFG RD | .015 | -- | -- |
| lIFG-lIPS FA | -.449 | -- | -- |
| rIFG-rIPS MD | .237 | -- | -- |
| rIFG-rIPS RD | .107 | -- | -- |
| lIPS-rIPS RD | .544** | .560** | 2.582** |
| Note: * p < .01, ** p < .05, *** p < .005 | | | |

| **Table S6.** Correlations for the ADHD group between absolute and relative motion with white matter structural metrics in order to ensure motion did not confound white matter structural metrics as it has been previously shown to do. | | | |
| --- | --- | --- | --- |
|  | N | Correlation Coefficient | p |
| lIFG-lIPS Volume |  |  |  |
| Relative Motion | 33 | -.050 | .784 |
| Absolute Motion | 33 | -.214 | .232 |
| rIFG-rIPS Volume |  |  |  |
| Absolute Motion | 33 | .136 | .449 |
| Relative Motion | 33 | .267 | .134 |
| IFG-IFG Volume |  |  |  |
| Relative Motion | 32 | .075 | .682 |
| Absolute Motion | 32 | .137 | .454 |
| IPS-IPS Volume |  |  |  |
| Absolute Motion | 32 | .181 | .321 |
| Relative Motion | 32 | .269 | .137 |
| lIFG-lIPS FA |  |  |  |
| Absolute Motion | 33 | .120 | .505 |
| Relative Motion | 33 | -.121 | .503 |
| rIFG-rIPS FA |  |  |  |
| Absolute Motion | 33 | -.012 | .949 |
| Relative Motion | 33 | -.335 | .057 |
| IFG-IFG FA |  |  |  |
| Absolute Motion | 32 | .090 | .625 |
| Relative Motion | 32 | -.201 | .269 |
| IPS-IPS FA |  |  |  |
| Absolute Motion | 32 | .151 | .410 |
| Relative Motion | 32 | .009 | .960 |
| lIFG-lIPS MD |  |  |  |
| Absolute Motion | 33 | .005 | .979 |
| Relative Motion | 33 | .185 | .303 |
| rIFG-rIPS MD |  |  |  |
| Absolute Motion | 33 | -.028 | .876 |
| Relative Motion | 33 | .166 | .355 |
| IFG-IFG MD |  |  |  |
| Absolute Motion | 32 | .335 | .061 |
| Relative Motion | 32 | .298 | .098 |
| IPS-IPS MD |  |  |  |
| Absolute Motion | 32 | .232 | .201 |
| Relative Motion | 32 | .278 | .124 |
| lIFG-lIPS AD |  |  |  |
| Absolute Motion | 33 | .058 | .749 |
| Relative Motion | 33 | .036 | .843 |
| rIFG-rIPS AD |  |  |  |
| Absolute Motion | 33 | -.014 | .821 |
| Relative Motion | 33 | -.198 | .269 |
| IFG-IFG AD |  |  |  |
| Absolute Motion | 32 | .317 | .077 |
| Relative Motion | 32 | .039 | .831 |
| IPS-IPS AD |  |  |  |
| Absolute Motion | 32 | .150 | .404 |
| Relative Motion | 32 | .152 | .399 |
| lIFG-lIPS RD |  |  |  |
| Absolute Motion | 33 | -.023 | .898 |
| Relative Motion | 33 | .181 | .312 |
| rIFG-rIPS RD |  |  |  |
| Absolute Motion | 33 | -.010 | .956 |
| Relative Motion | 33 | .348 | .047 |
| IFG-IFG RD |  |  |  |
| Absolute Motion | 32 | .075 | .678 |
| Relative Motion | 32 | .272 | .126 |
| IPS-IPS RD |  |  |  |
|  | 32 | .023 | .897 |
|  | 32 | .196 | .274 |
| Note: Correlations were tested using Spearman’s Rank Correlational Coefficient  Note: No correlations remained statistically significant following multiple comparison correction  Note: There is variation in the number of participants for each variable due to variation in the number of datasets lost for each variable due to outlier removal  Note: The P values reported are raw values, none of which survived multiple comparison correction | | | |

| **Table S7.** Correlations for the Controls group between absolute and relative motion with white matter structural metrics in order to ensure motion did not confound white matter structural metrics as it has been previously shown to do. | | | |
| --- | --- | --- | --- |
|  | N | Correlation Coefficient | P |
| lIFG-lIPS Volume |  |  |  |
| Relative Motion | 19 | -.190 | .436 |
| Absolute Motion | 19 | -.172 | .481 |
| rIFG-rIPS Volume |  |  |  |
| Absolute Motion | 19 | -.240 | .323 |
| Relative Motion | 19 | -.134 | .584 |
| IFG-IFG Volume |  |  |  |
| Relative Motion | 19 | .113 | .644 |
| Absolute Motion | 19 | .226 | .352 |
| IPS-IPS Volume |  |  |  |
| Absolute Motion | 18 | -.165 | .512 |
| Relative Motion | 18 | -.069 | .787 |
| lIFG-lIPS FA |  |  |  |
| Absolute Motion | 19 | -.347 | .145 |
| Relative Motion | 19 | -.026 | .917 |
| rIFG-rIPS FA |  |  |  |
| Absolute Motion | 19 | -.067 | .786 |
| Relative Motion | 19 | .158 | .581 |
| IFG-IFG FA |  |  |  |
| Absolute Motion | 19 | -.283 | .240 |
| Relative Motion | 19 | -.047 | .849 |
| IPS-IPS FA |  |  |  |
| Absolute Motion | 18 | .235 | .349 |
| Relative Motion | 18 | .296 | .233 |
| lIFG-lIPS MD |  |  |  |
| Absolute Motion | 19 | .147 | .549 |
| Relative Motion | 19 | -.122 | .619 |
| rIFG-rIPS MD |  |  |  |
| Absolute Motion | 19 | .054 | .827 |
| Relative Motion | 19 | .049 | .841 |
| IFG-IFG MD |  |  |  |
| Absolute Motion | 19 | .622 | .006 |
| Relative Motion | 19 | .147 | .561 |
| IPS-IPS MD |  |  |  |
| Absolute Motion | 18 | .029 | .911 |
| Relative Motion | 18 | -.464 | .061 |
| lIFG-lIPS AD |  |  |  |
| Absolute Motion | 19 | -.006 | .980 |
| Relative Motion | 19 | -.225 | .354 |
| rIFG-rIPS AD |  |  |  |
| Absolute Motion | 19 | -.224 | .357 |
| Relative Motion | 19 | -.064 | .794 |
| IFG-IFG AD |  |  |  |
| Absolute Motion | 19 | -.206 | .412 |
| Relative Motion | 19 | -.338 | .170 |
| IPS-IPS AD |  |  |  |
| Absolute Motion | 18 | -.127 | .615 |
| Relative Motion | 18 | -.555 | .017 |
| lIFG-lIPS RD |  |  |  |
| Absolute Motion | 19 | .328 | .170 |
| Relative Motion | 19 | .037 | .880 |
| rIFG-rIPS RD |  |  |  |
| Absolute Motion | 19 | .168 | .492 |
| Relative Motion | 19 | .006 | .980 |
| IFG-IFG RD |  |  |  |
| Absolute Motion | 19 | .583 | .009 |
| Relative Motion | 19 | .255 | .292 |
| IPS-IPS RD |  |  |  |
|  | 18 | .270 | .264 |
|  | 18 | .009 | .971 |
| Note: Correlations were tested using Spearman’s Rank Correlational Coefficient  Note: No correlations remained statistically significant following multiple comparison correction  Note: There is variation in the number of participants for each variable due to variation in the number of datasets lost for each variable due to outlier removal  Note: The P values reported are raw values, none of which survived multiple comparison correction | | | |
